# Dopamine Modulation of Spike-Timing-Dependent Plasticity for Spatio-Temporal Spike Pattern Detection in Single Neurons

**DOI:** 10.64898/2025.12.28.696794

**Authors:** Ayaka Kotajima, Shunta Furuichi, Takashi Kohno

## Abstract

Dopamine (DA) is an important neuromodulator that has been suggested to play key roles in a range of neuropsychiatric disorders. However, its computational impact at the neural circuit level has not been elucidated. Here, we extended a spike-timing-dependent plasticity (STDP)-based single-neuron spatio-temporal pattern detection model by incorporating a DA input and DA-type STDP modulation. We analyzed the direct effect of the DA-type STDP curve and the effect of DA release. We found that the DhA-type STDP curve accelerates learning but may promote coarse potentiation of the synaptic efficacy, which leads to false detections. It was also shown that DA can improve the pattern-detection performance up to a 44 % success rate (2.44 × control) with no false detection, but a DA release with an excessive amount or with too short a delay induces false detection. These results suggest that DA can maintain the pattern detection ability only when released within a limited concentration and a sufficient delay.

## 1 Introduction

Dopamine (DA) is one of the major neuromodulators in the central nervous system. By acting on widespread receptors and modulating synaptic plasticity and neuronal excitability, DA influences key cognitive functions such as attention, learning, and decision-making [1]. Dysregulation of dopaminergic signaling has also been implicated in a range of neuropsychiatric conditions, including attention-deficit/hyperactivity disorder (ADHD), autism spectrum disorder (ASD), and schizophrenia [2]. Importantly, these disorders place a substantial burden on individuals and society. Approximately 1/8 of the world’s population is afflicted [3], and current treatments still have major limitations. For example, while pharmacological interventions can be effective for some symptom domains, cognitive impairments in schizophrenia remain difficult to treat and continue to be a central obstacle to functional recovery [4].

A large body of prior works has investigated how DA relates to brain function and dysfunction at macroscopic and clinical levels (e.g., [5–8]). These studies have established important associations between dopaminergic signaling and macroscopic neural activity. However, explaining the mechanistic principles of the DA’s effects by such associations remains challenging. Because the brain is an extremely complex dynamical system, a simple one-to-one mapping between stimulus and response does not elucidate the mechanisms of the DA’s pharmacological effects at the neural circuit level.

On the other hand, the neural circuit level analyses are still limited. To the best of our knowledge, there is little preceding work other than the study by Wert-Carvajal et al. [9]. They studied an abstracted hippocampal network using a spiking neural network (SNN) model and analyzed the effects of DA- and 5-HT-type spike-timing-dependent plasticity (STDP) curves, demonstrating their impact at the level of the abstracted hippocampal circuit. The effect of DA on more fundamental single neurons’ information processing has not yet been analyzed.

In this study, we investigated the effect of DA on a single neuron by extending a spatio-temporal spike pattern detection model [10] based on STDP [11, 12]. A DA input was incorporated into the single-neuron model by Masquelier et al. [10]. Here, the effect of DA was modeled by a modified STDP learning curve reported by Zhang et al. [13] (DA-type STDP curve). We evaluated how the DA-type STDP curve modulates the performance of the spike pattern detection task with two setups. The first aimed to evaluate the direct effect of the DA-type STDP curve, and the second aimed to evaluate the effect of DA release under a more biologically realistic condition.

By focusing on a single neuron, we aim to provide a foundation for a mechanistic and bottom-up perspective on the dopaminergic modulation of neural circuits’ computation. This work is intended as a first step toward bridging macroscopic clinical observations and circuit-level principles, ultimately contributing to a clearer understanding of how DA can contribute to cortical information processing through synaptic plasticity.

## 2 Methods

### 2.1 Model

We incorporated a DA input into the spatio-temporal spike pattern detection model by Masquelier et al. [10]. The spiking dynamics are composed of a leaky integrate-and-fire (LIF) spiking mechanism with a double-exponential synapse model. In the study by Masquelier et al. [10], an asymmetric STDP was shown to enable a neuron to detect a spatio-temporal spike pattern embedded in noise (Poisson spikes). We adopted this model among single-neuron learning models because it provides one of the best balances between biological plausibility and computational cost.

The model equations are as follows. Let the membrane potential be

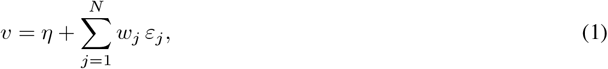

where *ε*_*j*_ is the excitatory postsynaptic potential (EPSP) raised by the *j*-th synapse, *η* is a variable generating spiking activity, and *w*_*j*_ is the synaptic weight of the *j*-th synapse. The total number of synaptic inputs *N* is 2000. Variables *η* and *ε*_*j*_ follow

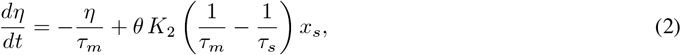

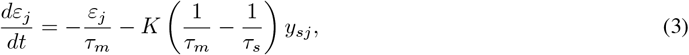

where *θ* = 500 is the threshold, *τ*_*m*_ = 10 ms is the membrane time constant, *τ*_*s*_ = 2.5 ms is the synaptic time constant, and *K* = 2.12, *K*_2_ = 4 are constants. The variables on the right-hand side of the above equations, *x*_*s*_ and *y*_*sj*_, evolve as

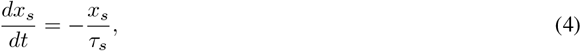

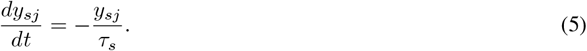

When *v* exceeds the threshold, the variables are reset to *η* = 2*θ, ε*_*j*_ = 0, and *x*_*s*_ = 1. At an arrival time *t*_*j*_ of a presynaptic spike on the *j*-th synapse, *y*_*sj*_ is reset to 1.

Synaptic weights *w*_*j*_ are updated by an asymmetric STDP rule. The change Δ*w*_*j*_ of *w*_*j*_ depends on the time difference between the presynaptic spike arrival time at synapse *j, t*_pre*j*_, and the neuron’s spike time *t*_post_. The baseline function is shown as Ctrl (blue) in Figure 1. Based on the experimental findings of Zhang et al. [13] on how DA modulates the STDP curve, we modeled the STDP curve under DA (orange).

**Figure 1:**
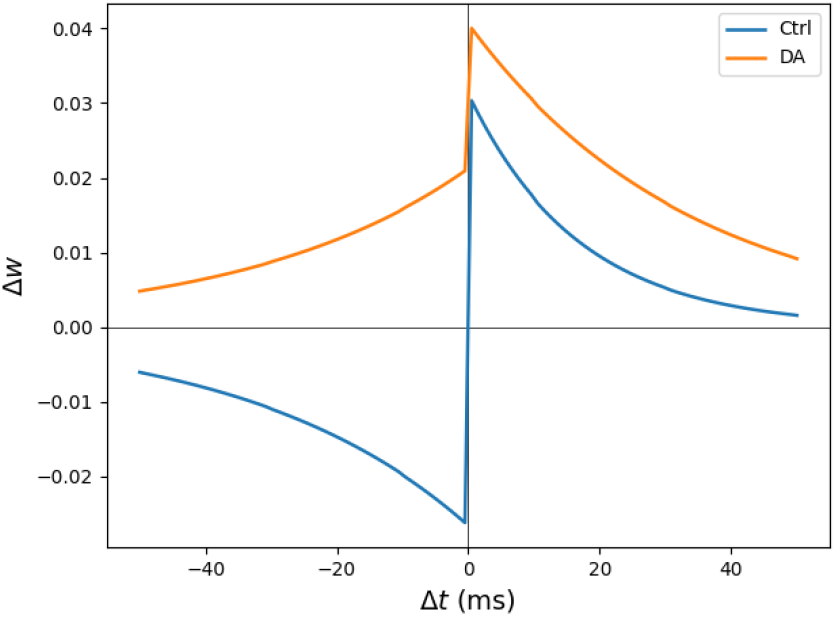
spike-timing-dependent plasticity (STDP) curves without DA (blue plot) and with Dopamine (DA) (orange plot). An asymmetric STDP curve was assumed for the control condition (without DA), while an all-positive curve was assumed under the existence of DA [13].

The STDP curve is

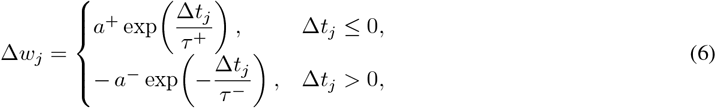

where Δ*t*_*j*_ = *t*_post_ − *t*_pre*j*_, *τ* ^+^, *τ* ^−^ are time constants, and *a*^+^, *a*^−^ are amplitude constants. For the baseline, we used *τ* ^+^ = 16.8 ms, *τ* ^−^ = 33.7 ms, *a*^+^ = 0.03125, *a*^−^ = 0.85 × 0.03125 (same as [10]). Under DA, 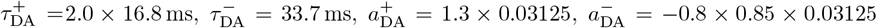 . Following learning, the weights were clipped to [0, 1].

### 2.2 Simulation setups

#### 2.2.1 (i) Switched STDP curve model

First, to investigate the effects of the DA-type learning curve, we switched between the blue and orange curves in Fig. 1.

The input spike trains were generated by the same procedure as in the study by Masquelier et al. [10]. Independent Poisson spike trains with rates 0 Hz to 90 Hz for 2000 synapses were generated. The first 50 ms spatio-temporal pattern was selected as the detection target, which was embedded with the rate of 10 % (approximately twice per second). To reflect the spike timing fluctuation, a Gaussian jitter with a mean of 0 ms and standard deviation (SD) of 1 ms to 5 ms (1 ms step) was applied. Then, 10 Hz Poisson spikes were added to the spike trains.

The DA-type STDP curve was applied for all or a part of the 50 ms target spatio-temporal patterns. In case of the partial application, the detection target pattern (50 ms) was partitioned into five 10 ms slots (1–5). In a designated slot, the STDP curve was switched to the DA-type with a probability of 10 % to 100 % in each simulation run. Because the effect of the DA-type STDP curve is very strong, as shown in the results section, we tested whether narrowing the time width and reducing the amplitude of the DA-type STDP curve could mitigate its effect. We uniformly scaled 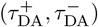 by *r*_*τ*_ ∈ {0.1, 0.2, …, 1.0} and 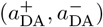 by *r*_*a*_ ∈ {0.1, 0.2, …, 1.0}.

In addition to the DA application slot and the STDP curve scaling (*r*_*τ*_, *r*_*a*_), we examined the effects of the initial synaptic weight *w*_0_ ∈{0.30, 0.35, 0.40} and the simulation time *T* ∈{50 s, 100 s, 200 s} . For each condition and each jitter level (1–5 ms), we simulated 50 independent spike trains with different random seeds (50 simulation runs for each parameter setting). A run was considered successful if, within the last 20 s, the detection rate was ≥96 % with zero false alarms.

#### 2.2.2 (ii) Exponentially decaying DA model

This setup aims to investigate the effect of DA in more realistic conditions. Here, we considered the dynamics of DA in brain tissue by incorporating its time constant. Considering that the time constant is about 400 ms [14], the appearance rate of the target 50 ms pattern was reduced to 1 % (approximately once per 5 s). In this setup, the spike-time jitter was fixed at 1 ms. The transmission delay of the DA pathway was also included in the model. Specifically, DA was assumed to be released with a fixed delay *δ* after the target pattern appeared, with a probability of 10 % to 100 %. At the release of DA, the concentration *m* is reset to *m*_0_ and then decays exponentially according to the following equation.

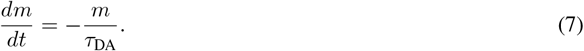

Here, *δ* ∈ {0 ms, 10 ms, 20 ms, 30 ms}, *τ*_DA_ = 400 ms, and *m*_0_ ∈{0.01, 0.02, …, 0.05, 0.10, …, 0.50} . Note that *m* is a normalized variable that indicates the rate of effectiveness of the DA. When *m* = 1.0, the STDP curve completely matches the DA-type one (the orange curve in Figure 1). The effective weight update under DA was a weighted average:

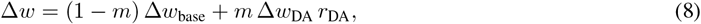

where Δ*w*_base_ and Δ*w*_DA_ are Δ*w* at the control condition and under DA, respectively. Parameter *r*_DA_ represents the efficacy of DA on the plasticity mechanism. A smaller value is selected for this scaling parameter to simulate a milder DA effect. We systematically varied *r*_DA_ ∈{0.1, 0.2, …, 1.0} and initial concentration *m*_0_ ∈{0.03, 0.05, 0.10, 0.15, …, 0.50} to examine its effect on synaptic modification. The simulation time was 200 s with an initial weight of *w*_0_ = 0.30. We used the last 50 s for the evaluation. The input spike trains and the detection success criterion were the same as the previous setup.

## 3 Results

### 3.1 (i) Switched STDP curve model

We first examined the direct effect of the DA-type STDP curve on pattern detection using the switched STDP curve model.

For each parameter condition, we executed 50 simulation runs with independent Poisson input spike trains with different random seeds. For each trial, we evaluated the result using the last 20 s of the simulation data. A run was counted as successful when the detection rate in this evaluation window was at least 96 % and no spike occurred outside the detection target patterns. Here, the detection rate is the rate of the bins of the detection target pattern with a spike divided by the total number of bins of the detection target pattern. In addition, we quantified the false alarm fraction, defined as the proportion of runs that satisfied the detection criterion (detection rate ≥96 %) but produced at least one false alarm. Simulations were implemented in Brian2 [15] using explicit Euler integration (Δ*t* = 0.1 ms). Random seeds were set as *S*_0_ = 1000 and *S*_*i*_ = *S*_0_ + *i* (*i* = 0, …, 49).

When the DA-type STDP curve was applied all over the detection target pattern, the model never reached the success criterion: the success rate was 0 % and all runs exhibited false alarms (100 % false alarm fraction), indicating that uniform DA modulation makes the neuron excessively sensitive. Therefore, we applied DA-type STDP for only one 10 ms slot with a release probability (see Sec. 2.2.1).

Figure 2 shows the results with the baseline condition (*w*_0_ = 0.3, *r*_*τ*_ = *r*_*a*_ = 1.0, and *T* =100 s). Each point represents the success rate, the proportion of successful runs among 50 runs. Here, error bars indicate 95 % Wilson confidence intervals for the binomial proportion. When the jitter noise was moderate (2 ms), the moderate DA release probability (40 %) improved the detection success rate compared to the control condition without DA (the value at the release probability 0 %). However, further increase of the release probability suppressed the success rate. This deterioration was accompanied by a marked increase in the false alarm fraction (see Figure 2(b)). The impact of DA was more pronounced for larger jitter values, but the success rate was lower across the release probability. While DA could enhance the success rate, it also facilitated potentiation for weakly related inputs and therefore made the neuron more prone to false detections, especially when the release probability was higher.

**Figure 2:**
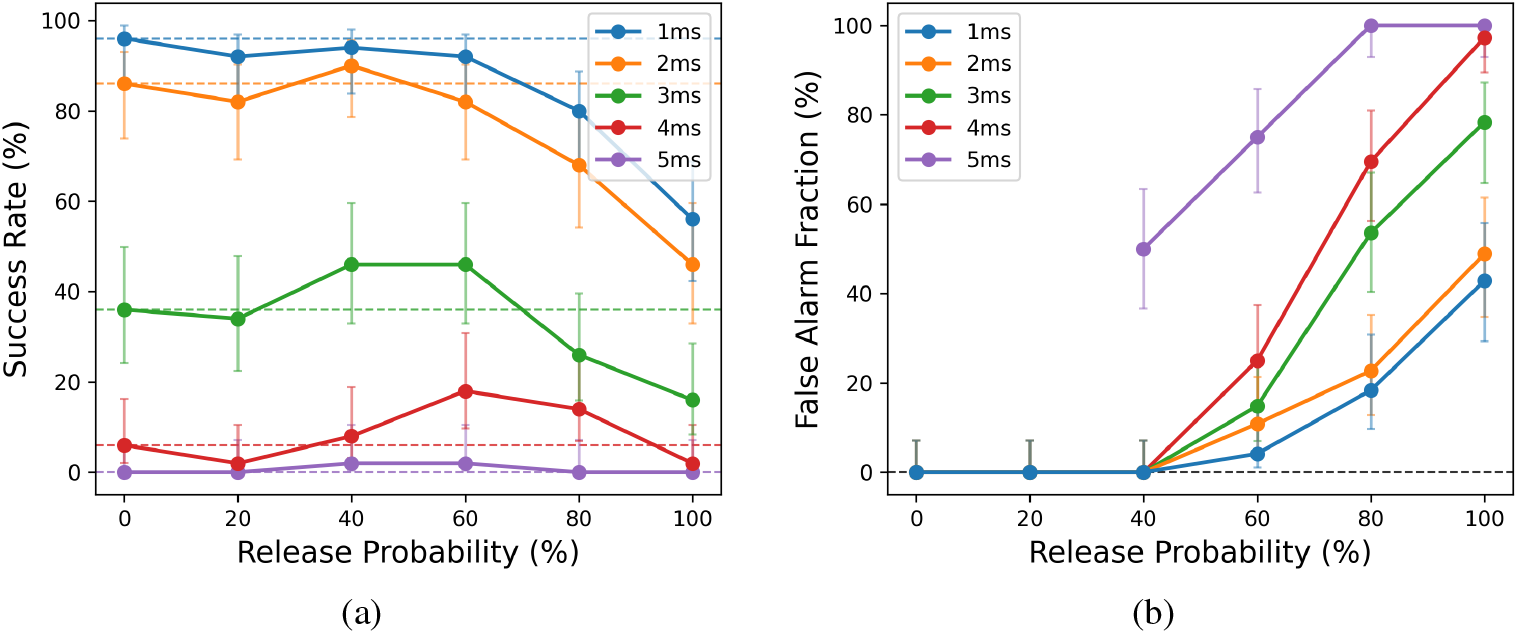
Dependence of (a) the detection success rate and (b) the false alarm fraction on dopamine (DA) release probability and spike-time jitter under the baseline condition: the initial synaptic weight was *w*_0_ = 0.30, *r*_*τ*_ = *r*_*a*_ = 1.0, DA was applied only in the third 10 ms slot of the 50 ms embedded pattern, and the total simulation time was 100 s. Each point represents the average over 50 runs, and the error bars indicate 95 % Wilson confidence intervals for the binomial proportion (Newcombe–Wilson method).

To investigate the success rate enhancement effect for moderate jitter noise (2 ms), we first tested the dependence on the learning duration *T* (see Figure 3(a)). For a shorter learning duration (*T* = 50 s), moderate DA release (20–40%) led to a clear gain in success rate, whereas for a longer learning duration (*T* = 200 s) this benefit was reduced, especially when the release probability was higher, because incorrect synapses had more opportunities to be strengthened. These results suggest that the DA-type STDP curve helps quick learning by widening the LTP time window. Next, we visualized the dependence of the false alarm fraction on the release timing slot (Figure 3(b)). Here, the release probability was 60 %. False alarms were particularly frequent when DA was released in the first slot of the pattern. In this case, DA was applied immediately after a random, non-pattern segment that would normally induce LTD. The presence of DA converted these random presynaptic spikes into LTP, which reinforced irrelevant synapses. Finally, the dependence on the initial synaptic weight *w*_0_ was evaluated (Figure 4). Higher initial weights, corresponding to higher intrinsic excitability of the neurons, led to more frequent false alarms and diminished the beneficial effect of DA.

**Figure 3:**
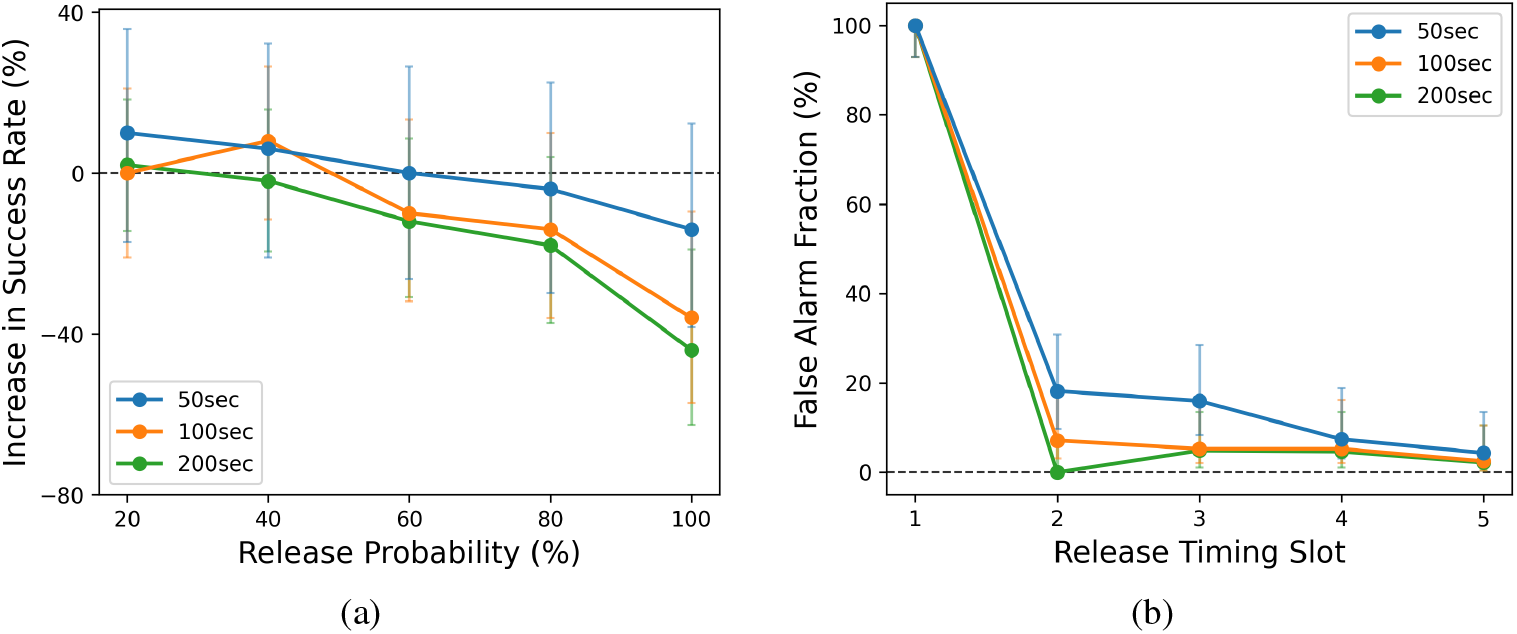
Dependence of the effect of dopamine (DA) on the learning duration *T* for jitter noise 2 ms (other parameters are as in Fig. 2). (a) Increase in detection success rate relative to the control condition without DA; each point represents the average over *n* = 50 runs and the error bars denote 95 % Wilson confidence intervals for the difference of binomial proportions (Newcombe–Wilson method). The line of 0 % corresponds to the success rate for the release probability 0 % for each value of *T* . (b) False alarm fraction as a function of the DA release slot (horizontal axis) at a release probability of 60 %.

**Figure 4:**
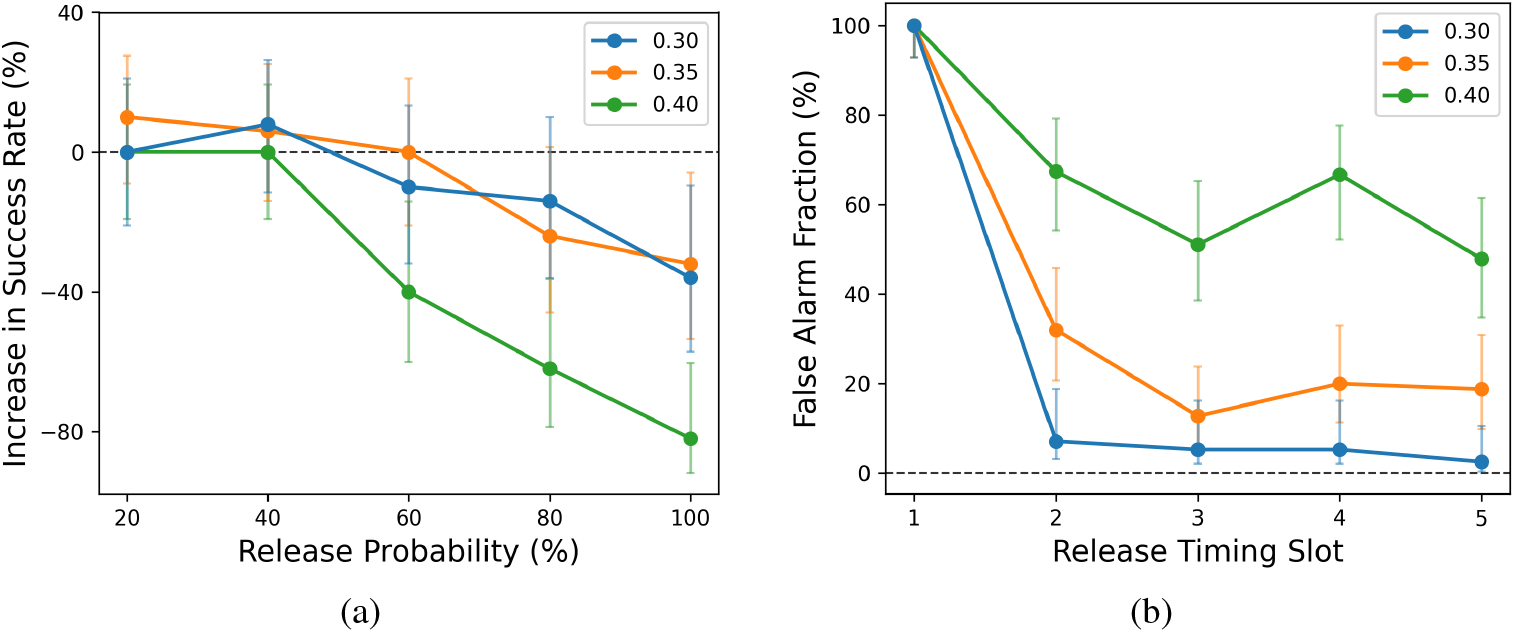
Dependence of the effect of dopamine (DA) on the initial synaptic weight *w*_0_ for jitter 2 ms (other parameters are as in Fig. 2). Each point represents the average over 50 runs, and the error bars indicate 95 % Wilson confidence intervals for the binomial proportion (Newcombe–Wilson method).

The above results indicate that the DA-type STDP curve has a trade-off between the beneficial (improving the success rate and accelerating the learning speed) and detrimental (increasing the false alarm fraction) effects. Here, we examined how this trade-off depends on the scale of the STDP curve. In this analysis, the time constant and amplitude of the DA-type STDP curve were multiplied by factors *r*_*τ*_ and *r*_*a*_, respectively. Figure 5 shows the dependence on (*r*_*a*_, *r*_*τ*_) for jitter 2 ms and DA release probability of 100 %. Even at such a high release probability, narrowing the STDP window substantially suppressed the false alarm fraction. For instance, under the baseline condition (*r*_*a*_, *r*_*τ*_) = (1.0, 1.0) the detection success rate was only 46 % and the false alarm fraction was comparably high (approximately 49 %), whereas scaling the DA-type STDP curve to (*r*_*a*_, *r*_*τ*_) = (0.4, 0.6) increased the success rate to 86 % and eliminated false alarms (0 %). The figures indicate that reducing *r*_*a*_ to half eliminates the false alarms and improves the success rate over 70 %.

**Figure 5:**
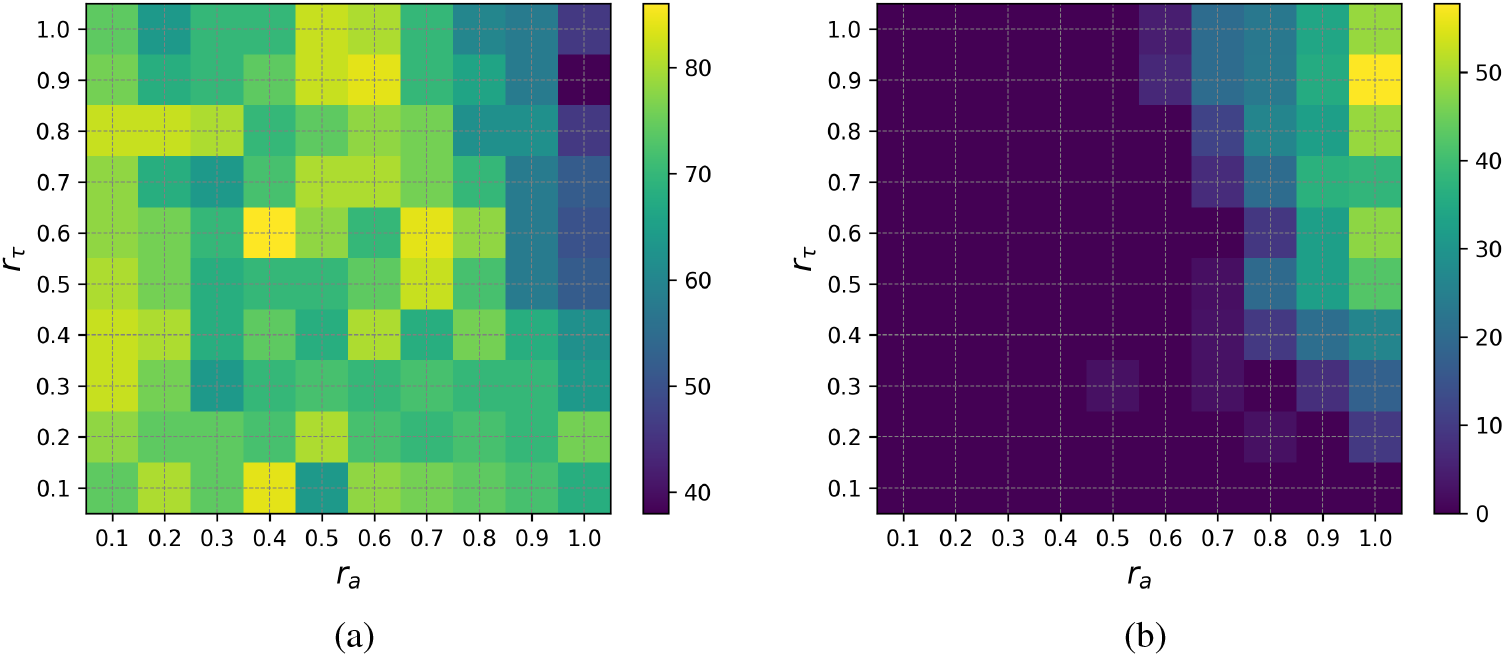
Effect of scaling the dopamine (DA)-type spike-timing-dependent plasticity (STDP) curve under the baseline condition and a DA release probability of 100 %. (a) Detection success rate and (b) false alarm fraction as functions of the scaling factors (*r*_*a*_, *r*_*τ*_). Each point indicates the average over 50 runs.

Overall, these results show that the DA-type STDP curve accelerates learning and enables rapid acquisition of the detection target pattern, but at the cost of “coarse” learning, the neuron tends to spike in response to non-target patterns.

When the DA release probability or the baseline excitability is too high, the neuron is more likely to learn incorrect patterns and to fire in non-target patterns. By scaling the DA-type STDP curve, we could suppress these false alarms, even at high DA release probabilities, because this manipulation selectively weakens potentiation for non-matching inputs while preserving strong potentiation at the correct pattern timing.

### 3.2 (ii) Exponentially decaying DA model

Next, we examined how DA can modulate the spatio-temporal pattern-detection performance. As described in Section 2.2.2, the embedded pattern appeared with a probability of 1 % and the DA concentration *m*(*t*) decayed exponentially.

In the control condition without DA, the neuron detected only about 18 % of the embedded patterns despite the long learning duration (*T* =200 s).

To highlight the effect of DA dynamics, we first fixed the scaling parameter to *r*_DA_ = 1.0 and investigated how the initial concentration *m*_0_ and DA release probability affect the success rate and false alarm fraction (Figure 6). When *m*_0_ was larger than 0.10, the success rate tended to drop as the release probability increased (Figure 6(a)), which was accompanied by the increase of the false alarm fraction (Figure 6(b)). In contrast, when *m*_0_ was 0.10 or smaller, the success rate tended to increase in response to the increase in the release probability. When *m*_0_ was 0.10, the model yielded the best performance: the detection success rate reached 44 % with no false alarms, 2.44 times higher than the 18 % in the control condition without DA. Although a success rate of 44 % at the single-neuron level may seem modest, it would provide a substantial gain when many neurons act as weak learners in parallel, suggesting that DA can enhance the ability to detect the events that occur less frequently in cortical circuits.

**Figure 6:**
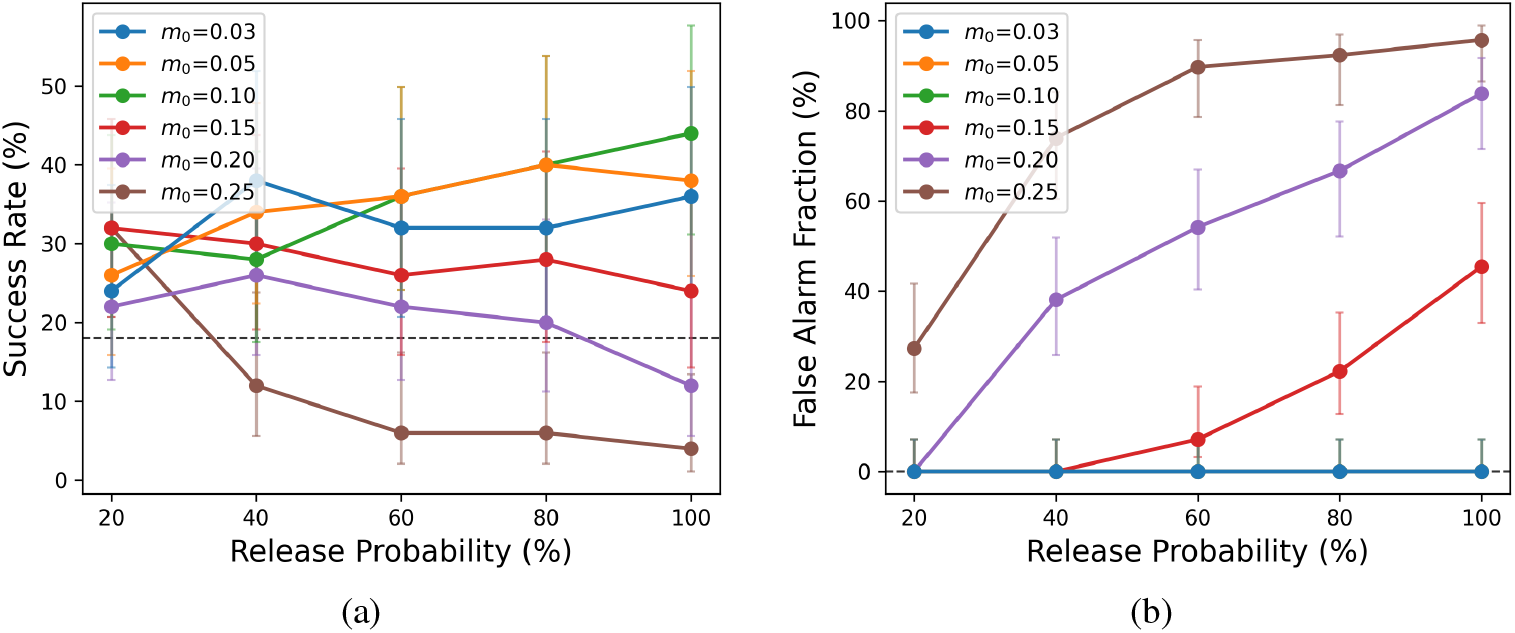
Dependence of (a) the detection success rate and (b) the false alarm fraction on initial dopamine (DA) concentration *m*_0_ and the DA release probability. Scaling parameter *r*_DA_ was fixed at 1.0 and delay *δ* was 10 ms. Each point represents the average over 50 runs, and the error bars indicate 95 % Wilson confidence intervals for the binomial proportion (Newcombe–Wilson method).

Second, we examined how the DA scaling parameter *r*_DA_ and delay *δ* in the DA pathway influence the detection performance. Figure 7 shows how the detection performance depended on the combination of *r*_DA_ and *m*_0_. In the figure, *δ* was 10 ms. Weakening *r*_DA_ suppressed false alarms even when *m*_0_ was high. However, the best results were obtained when *m*_0_ was relatively low (0.10) and *r*_DA_ was high (0.9–1.0). The success rate was 44 % without false alarms (the blue boxes in Figure 7(a)). When *r*_DA_ was lower, moderate *m*_0_ (0.3–0.8) yielded a higher success rate (48 %) but with 4 % to 8 % false alarm fraction (red boxes in the figure). In the figure, a high success rate tends to appear for larger *r*_DA_ when *m*_0_ is smaller. It is natural that there are best balance points between the initial concentration and the effectiveness of DA. However, the figure also indicates that a very low initial concentration yields relatively good performance regardless of *r*_DA_. Figure 8 shows the dependence on *δ*. When DA was released without delay (*δ* = 0 ms), the success rate was high only when *m*_0_ was very small, and was otherwise zero with frequent false alarms. Introducing a short delay of about *δ* = 10 ms markedly reduced the false alarm fraction while maintaining a high success rate, yielding the best overall performance (the blue boxes in Figure 8(a)). Further increasing the delay to *δ* ≥20 ms suppressed false alarms more effectively, but the improvement in success rate remained limited. Thus, to maximize the success rate under the strict condition of zero false alarms, a relatively short delay of around 10 ms was optimal.

**Figure 7:**
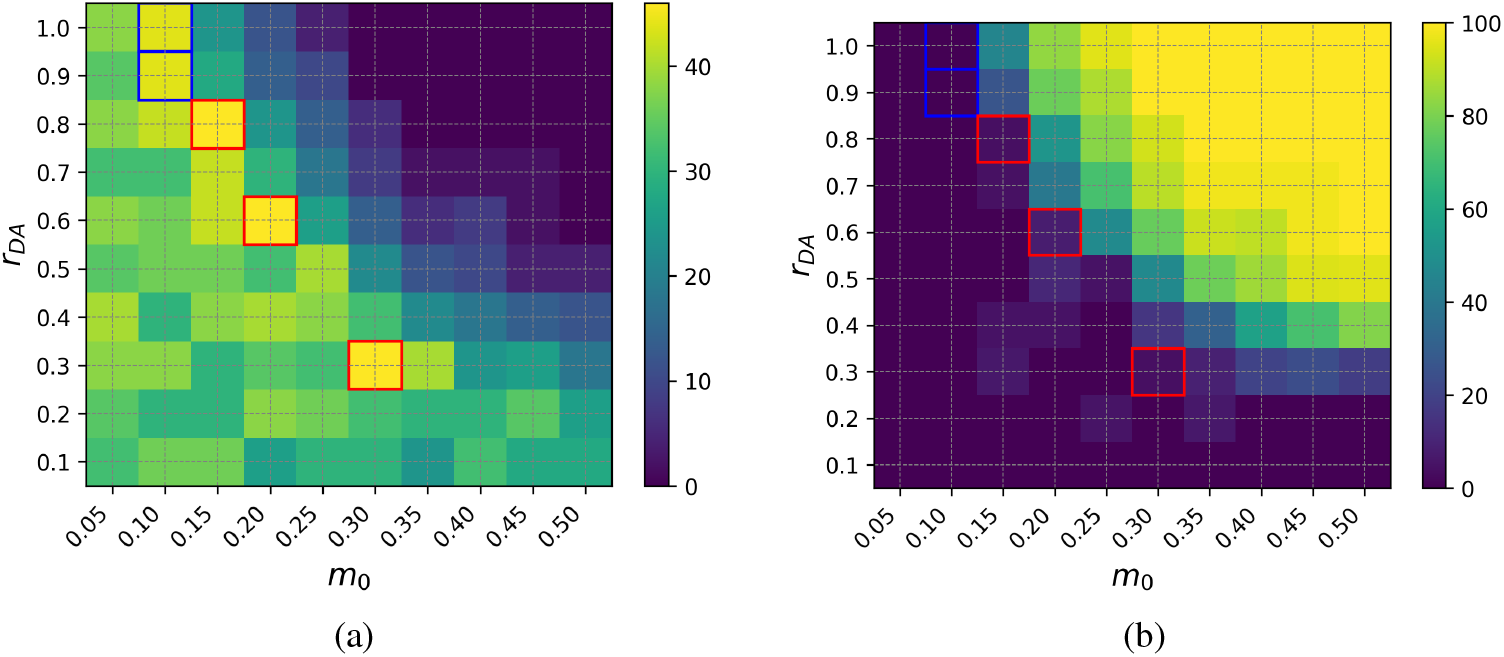
Dependence of the pattern-detection performance on initial dopamine (DA) concentration *m*_0_ and DA scaling parameter *r*_DA_. Panels show (a) the detection success rate and (b) the false alarm fraction as functions of *m*_0_ and *r*_DA_ for a fixed DA release delay *δ*. Delay *δ* was 10 ms and release probability was 100 %. Each point indicates the average over 50 runs. In both panels, the points enclosed by a blue box indicate the best success rate with no false alarms. The points with a red box have a higher success rate, but with false alarms.

**Figure 8:**
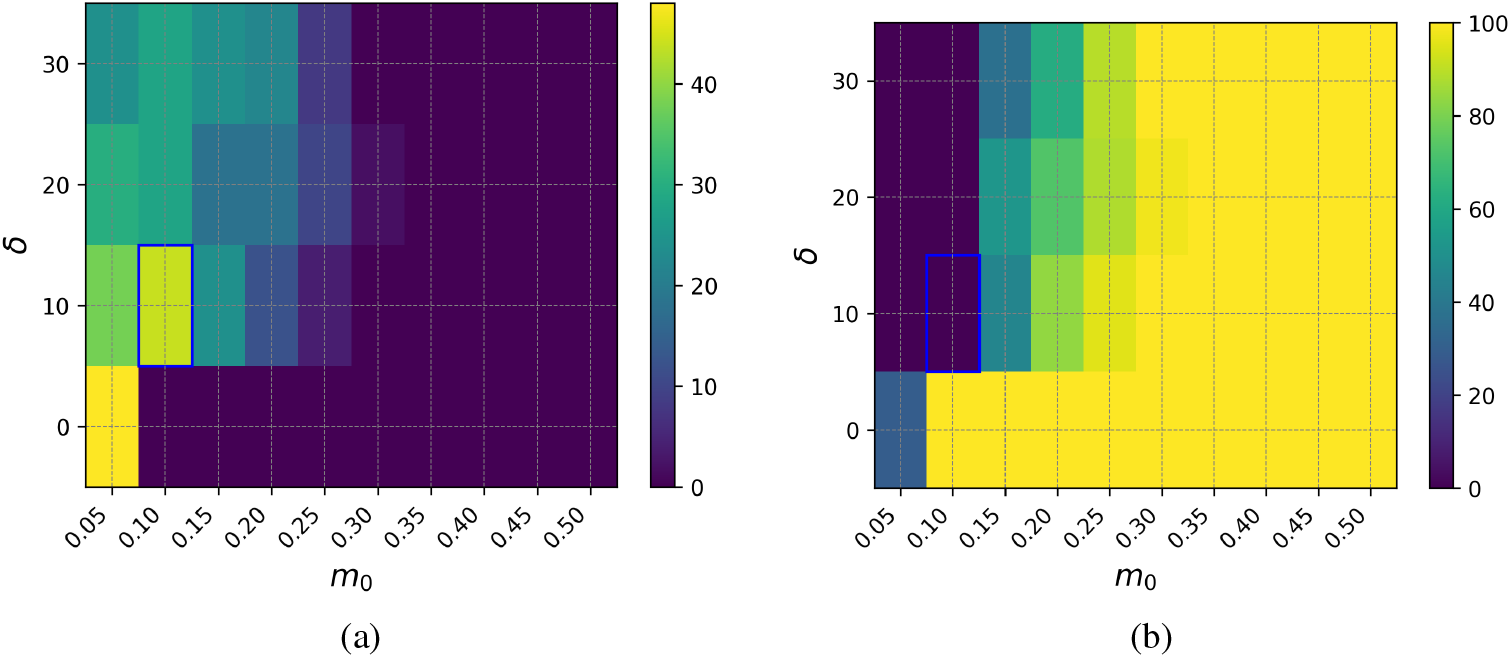
Dependence of pattern-detection performance on initial dopamine (DA) concentration *m*_0_ and DA release delay *δ*. The panels show (a) the detection success rate and (b) the false alarm fraction. Scaling parameter *r*_DA_ was fixed at 1.0 and the release probability was 100 %. Each point indicates the average over 50 runs. The point enclosed by a blue box indicates the best success rate with no false alarms.

Overall, these results indicate that the exponentially decaying DA model also exhibits a trade-off between enhanced pattern detection and robustness against false alarms. The pattern-detection performance is governed by the interaction between initial DA level *m*_0_ and the scaling parameter *r*_DA_. The best success rate without false alarm was achieved when *m*_0_ was low and *r*_DA_ was high. But when *r*_DA_ was lower, moderate *m*_0_ could achieve good results. Larger *m*_0_ or higher *r*_DA_ increased the likelihood of false detections. Conversely, false alarms were suppressed when the DA effect was weak, even at high *m*_0_, but did not yield the best improvement in the success rate. Thus, an appropriate balance between the initial DA concentration and the “DA effect” represented by *r*_DA_ is suggested to be essential for reliable and accurate pattern detection. The importance of a short delay was also suggested.

## 4 Conclusion

In this study, we conducted a bottom-up analysis to clarify how DA affects neural computation at the circuit level by focusing on a single neuron, the most fundamental unit of neural circuits. By incorporating a DA input into the spatio-temporal spike pattern detection model of Masquelier et al. [10] and modeling DA effects through the DA-type STDP learning rule reported by Zhang et al. [13], we investigated the effect of DA on a single neuron’s information processing.

Across two simulation setups, we consistently found that DA modulation introduces a trade-off between sensitivity (rapid learning and higher detection success) and specificity (robustness against false alarms). In the switched curve setup, the DA-type STDP curve was shown to accelerate learning and enable rapid acquisition of the target pattern, but also had a tendency to promote coarse learning that increased the number of false alarms. In the exponentially decaying DA setup, performance depended on the interaction among the initial DA concentration *m*_0_, the DA scaling parameter *r*_DA_, and the release delay *δ*. It was suggested that a lower initial DA concentration and a DA release delay of 10 ms or longer are advantageous for spatio-temporal spike pattern detection capability. The success rate reached 44 % (2.44 times the control condition without DA) with zero false alarms, indicating that DA can substantially enhance detection of infrequent events even at the single-neuron level when its magnitude and timing are appropriately balanced. It was also suggested that suppressing the initial DA concentration can effectively eliminate the false alarms without critically affecting the success rate.

These findings provide a bottom-up perspective: dopaminergic modulation can maintain good learning and detection of behaviorally relevant patterns, but excessive DA effect or mistimed DA signals may increase false detections by reinforcing non-target inputs. This sensitivity–specificity trade-off suggests that not only the amount of DA, but also its temporal alignment with synaptic events is critical for reliable information processing through plasticity. The relative insensitivity of the success rate to DA concentration compared to false detection suggests that suppressing the amount of DA release may be a good strategy to suppress false detection without critically damaging learning ability.

Future work should extend this framework beyond a single neuron to multiple neurons. It will be important to incorporate more biologically detailed DA dynamics and other neuromodulators such as 5-HT. These extensions may help bridge circuit-level principles with the cognitive impairments associated with neuropsychiatric disorders.

## Acknowledgements

This work was partly supported by Japan Science and Technology Agency (JST) Science and Technology Challenge Program for Next Generation, and UTokyoGSC-Next.

## References

[1] Ethan S. Bromberg-Martin, Masayuki Matsumoto, and Okihide Hikosaka. Dopamine in motivational control: rewarding, aversive, and alerting. Neuron, 68(5):815–834, 2010. doi:10.1016/j.neuron.2010.11.022.

[2] Yunyun Cai, Lingyan Xing, Tuo Yang, Rui Chai, Jiaqi Wang, Jingyin Bao, Weixing Shen, Sujun Ding, and Gang Chen. The neurodevelopmental role of dopaminergic signaling in neurological disorders. Neuroscience Letters, 741:135540, 2021. doi:10.1016/j.neulet.2020.135540.

[3] GBD 2019 Mental Disorders Collaborators. Global, regional, and national burden of 12 mental disorders in 204 countries and territories, 1990–2019: a systematic analysis for the global burden of disease study 2019. The Lancet Psychiatry, 9(2):137–150, 2022. doi:10.1016/S2215-0366(21)00395-3.

[4] Robert A. McCutcheon, Richard S. E. Keefe, and Philip K. McGuire. Cognitive impairment in schizophrenia: aetiology, pathophysiology, and treatment. Molecular Psychiatry, 28(5):1902–1918, 2023. doi:10.1038/s41380-023-01949-9.

[5] Robert A. McCutcheon, Anissa Abi-Dargham, and Oliver D. Howes. Schizophrenia, dopamine and the striatum: from biology to symptoms. Trends in Neurosciences, 42(3):205–220, 2019. doi:10.1016/j.tins.2018.12.004.

[6] Antoine Rogeau, Anne Jetske Boer, Eric Guedj, Arianna Sala, Iris E. Sommer, Mattia Veronese, Monique van der Weijden-Germann, Francesco Fraioli, and EANM Neuroimaging Committee. EANM perspective on clinical PET and SPECT imaging in schizophrenia-spectrum disorders: a systematic review of longitudinal studies. European Journal of Nuclear Medicine and Molecular Imaging, 52(3):876–899, 2025. doi:10.1007/s00259-024-06987-1.

[7] Hayley J. MacDonald, Rune Kleppe, Peter D. Szigetvari, and Jan Haavik. The dopamine hypothesis for ADHD: an evaluation of evidence accumulated from human studies and animal models. Frontiers in Psychiatry, 15:1492126, 2024. doi:10.3389/fpsyt.2024.1492126.

[8] Denis Paval. A dopamine hypothesis of autism spectrum disorder. Developmental Neuroscience, 39(5):355–360, 2017. doi:10.1159/000478725.

[9] Carlos Wert-Carvajal, Melissa Reneaux, Tatjana Tchumatchenko, and Claudia Clopath. Dopamine and serotonin interplay for valence-based spatial learning. Cell Reports, 39(2):110645, 2022. doi:10.1016/j.celrep.2022.110645.

[10] Timothée Masquelier, Rudy Guyonneau, and Simon J. Thorpe. Spike timing dependent plasticity finds the start of repeating patterns in continuous spike trains. PLOS ONE, 3(1):e1377, 2008. doi:10.1371/journal.pone.0001377.

[11] Henry Markram, Joachim Lübke, Michael Frotscher, and Bert Sakmann. Regulation of synaptic efficacy by coincidence of postsynaptic APs and EPSPs. Science, 275(5297):213–215, 1997. doi:10.1126/science.275.5297.213.

[12] Guo-qiang Bi and Mu-ming Poo. Synaptic modifications in cultured hippocampal neurons: dependence on spike timing, synaptic strength, and postsynaptic cell type. Journal of Neuroscience, 18(24):10464–10472, 1998. doi:10.1523/JNEUROSCI.18-24-10464.1998.

[13] Ji-Chuan Zhang, Pak-Ming Lau, and Guo-Qiang Bi. Gain in sensitivity and loss in temporal contrast of STDP by dopaminergic modulation at hippocampal synapses. Proceedings of the National Academy of Sciences of the United States of America, 106(31):13028–13033, 2009. doi:10.1073/pnas.0900546106.

[14] Abraham G. Beyene, Kristen Delevich, Jackson Travis Del Bonis-O’Donnell, David J. Piekarski, Wan Chen Lin, A. Wren Thomas, Sarah J. Yang, Polina Kosillo, Darwin Yang, George S. Prounis, Linda Wilbrecht, and Markita P. Landry. Imaging striatal dopamine release using a nongenetically encoded near infrared fluorescent catecholamine nanosensor. Science Advances, 5(7):eaaw3108, 2019. doi:10.1126/sciadv.aaw3108.

[15] Marcel Stimberg, Romain Brette, and Dan F. M. Goodman. Brian 2, an intuitive and efficient neural simulator. eLife, 8:e47314, 2019. doi:10.7554/eLife.47314.

